# Tonic and transient oscillatory brain activity during acute exercise

**DOI:** 10.1101/201749

**Authors:** Luis F. Ciria, Antonio Luque-Casado, Daniel Sanabria, Darias Holgado, Plamen Ch. Ivanov, Pandelis Perakakis

## Abstract

The physiological changes that occur in the main body systems and organs during physical exercise are well described in the literature. Despite the key role of brain in processing afferent and efferent information from organ systems to coordinate and optimize their functioning, little is known about how the brain works during exercise. The present study investigated tonic and transient oscillatory brain activity during a single bout of aerobic exercise. Twenty young males (19-32 years old) were recruited for two experimental sessions on separate days. Electroencephalographic (EEG) activity was recorded during a session of cycling at 80% (moderate-to-high intensity) of VO_2max_ (maximum aerobic capacity) while performing an oddball task where participants had to detect infrequent targets presented among frequent non-targets. This was compared to a (baseline) light intensity session (30% VO_2max_). The light intensity session was included to control for any potential effect of dual-tasking (i.e., pedaling and performing the oddball task). A warm-up and cool down periods were completed before and after exercise, respectively. A cluster-based nonparametric permutations test showed an increase in power across the entire frequency spectrum during the moderate-to-high intensity exercise, with respect to light intensity. Further, we found that the more salient target lead to lower increase in (stimulus-evoked) theta power in the 80% VO_2max_ with respect to the light intensity condition. On the contrary, higher decrease alpha and lower beta power was found for standard trials in the moderate-to-high exercise condition than in the light exercise condition. The present study unveils, for the first time, a complex brain activity pattern during acute exercise (at 80% of maximum aerobic capacity). These findings might help to elucidate the nature of changes that occur in the brain during physical exertion.

## Introduction

The dynamics and regulatory mechanism of body systems and organs such as muscles, joints, heart, lungs, etc. under physical exercise is well described in the literature [1][2][3][4][5][6]. However, little is known about how the brain works when exercising [7][8]. Here, we provide novel evidence on oscillatory brain activity during a single bout of aerobic exercise as a starting point to better understand the highly complex way in which the brain functions during physical exertion.

The study of brain activity in motion entails several methodological and technical issues (e.g. sweating, body movements, muscle potentials, etc.). This is probably the main reason why only a reduced number of studies have investigated brain activity during exercise, primarily using electroencephalography (EEG). The majority of these studies focused on changes in the alpha frequency band at frontal localizations [9] reporting increased alpha activity during exercise [10][11]. The only meta-analysis to date [12], however, indicates that alpha brain rhythm activation is not selective and maybe parallel by an increase of other brain rhythms, suggesting a possible power increase across the entire frequency spectrum during exercise. Notably, these early studies focused on the averaged steady state spectral activation across the entire exercise period and have not investigated transient modulation in brain rhythms in response to task relevant stimuli under physical effort.

An alternative line of investigations have focused on target-locked brain responses during exercise by means of EEG event-related brain potentials (ERPs) as a way to pinpoint the brain correlates of (task relevant) stimulus processing under physical effort [13][14][15][16][17][18]. Here, we take a step further by analysing power spectral changes time-locked to the (task) stimulus instead of ERPs. The event-related spectral perturbation (ERSP) analysis refers to transient decreases or increases in oscillatory brain activity locked to an event, which are thought to reflect the state of synchrony in a population of neurons [19][20][21], and it may provide complementary information to that of the ERPs. The study of ERSP during exercise was achieved by including an oddball task whereby participants had to detect infrequent targets presented among a sequence of frequent distractors (see Yagi et al. [18], for a similar approach although using ERPs).

Our participants exercised at two different intensities, corresponding to the 80% and 30% of their VO_2max_ (i.e., maximal aerobic capacity). This selection was motivated by previous evidence pointing to moderate-to-high acute exercise (between the 60% and 80% VO_2max_) as the key intensity to induce cognitive enhancement [22][23][24]. The 30% condition was included as the low-intensity exercise baseline (instead of a rest non-exercise condition) to control for potential dual-tasking (i.e., participants were both exercising and performing a cognitive task).

In sum, the current empirical study focused on the tonic oscillatory brain activity and, for the first time, on the event-related brain oscillatory responses during two sessions of acute aerobic exercise at different intensities (moderate-to-high and light) while performing an oddball task. Importantly, taking into account the sparse literature about the brain dynamics during physical exercise we took an exploratory approach (cf. Wagenmakers et al. [25]), with a bottom-up methodology by employing a stepwise cluster-based analysis without prior assumptions on any frequency range or brain localization.

## Material and Methods

### Participants

We recruited 20 young males with a high level of aerobic fitness (age between 18-31 years old, average age 23.9 years old) from the University of Granada (Spain). All participants met the inclusion criteria of reporting at least 8 hours of cycling or triathlon training per week, normal or corrected to normal vision, reported no neurological, cardiovascular or musculoskeletal disorders and were taking no medication. Note that high-fit cyclists and triathletes were selected because they are capable of maintaining a pedalling cadence at moderate-to-high intensity during long periods of time. Furthermore, they are able to keep a fixed posture over time, which notably reduces EEG movement artifacts. Their fitness level was verified by an incremental effort test (see below). Participants were required to maintain a regular sleep-wake cycle for at least one day before each experimental session and to abstain from stimulating beverages or any intense physical activity 24 hours before each session. All subjects gave written informed consent before the study. The protocol was in accordance with both, the ethical guidelines of the University of Granada, and the Declaration of Helsinki.

### Apparatus and materials

All participants were fitted with a Polar RS800 CX monitor (Polar Electro Öy, Kempele, Finland) to record their heart rate (HR) during the incremental exercise test. We used a ViaSprint 150 P cycle ergometer (Ergoline GmbH, Germany) to induce physical effort and to obtain power values, and a JAEGER Master Screen gas analyser (CareFusion GmbH, Germany) to provide a measure of gas exchange during the effort test. Oddball stimuli were presented on a 21-inch BENQ screen maintaining a fixed distance of 100 cm between the head of participants and the center of the screen. E-Prime software (Psychology Software Tools, Pittsburgh, PA, USA) was used for stimulus presentation and behavioural data collection.

### Fitness Assessments

Participants came to the laboratory, at least one week before the first experimental session, to provide the informed consent, complete an anthropometric evaluation (height, weight and body mass index [BMI]) and to familiarize with the oddball task. Subsequently, they performed an incremental cycle-ergometer test to obtain their maximal oxygen consumption (VO_2max_) which was used in the following experimental sessions to adjust the exercise intensity individually. The incremental effort test started with a 3 minutes warm-up at 30 Watts (W), with the power output increasing 10 W every minute. Each participant set his preferred cadence (between 60-90 rpm · min^-1^) during the warm-up period and was asked to maintain this cadence during the entire protocol. The test began at 60 W and was followed by an incremental protocol of 30 W every 3 minutes. Each step of the incremental protocol consisted of 2 minutes of stabilized load and 1 minute of progressive load increase (5 W every 10 seconds). The oxygen uptake (VO_2_ ml • min^-1^ • kg^−1^), respiratory exchange ratio (RER; i.e., CO_2_ production • O_2_ consumption^−1^), relative power output (W • Kg^−1^) and heart rate (bpm) were continuously recorded throughout the test.

### Experimental sessions

Participants completed two counterbalanced experimental sessions of approximately 100 min each. To avoid possible fatigue and/or training effects, visits to the laboratory were scheduled on different days allowing a time interval of 48-72 hours between sessions. On each experimental session, after 10’ warm-up on a cycle-ergometer at a power load of 30% of their individual VO_2max_, participants performed an oddball task for 20’ while pedalling either at 30% (light intensity exercise session) or 80% (moderate-intensity exercise session) of their VO_2max_. Upon completion of the oddball task, a 10’ cool down period at 30% of intensity followed (see Table 1). Each participant set his preferred cadence (between 60-90 rpm · min^-1^) before the warm-up and was asked to maintain this cadence throughout the session in order to match conditions in terms of dual-task demands.

### Oddball task

The visual oddball task was based on that reported in Sawaki and Katayama [26]. The task consisted of a random presentation of three visual stimuli, consisting of a frequent small blue circle (approximately 1.15° x 1.15°), a rare big blue circle (approximately 1.30° x 1.30°), and a rare red square (approximately 2.00° x 2.00°). Small blue circles were considered as standard stimuli (non-target), while big blue circles and red squares were considered as target stimuli. Stimuli were displayed sequentially on the center of the screen on a black background. Each trial started with the presentation of a blank screen in a black background for 1200 ms. Then, the stimulus was presented at a random time interval (between 0 and 800 ms) during 150 ms. Participants were instructed to respond to both targets by pressing a button connected to the cycle-ergometer handlebar with the thumb of their dominant hand and to not respond when standard stimuli were shown. Participants were encouraged to respond as accurately as possible. The target stimuli were randomly presented in 20% of trials (10% of target 1, 10% target 2) and the nontarget stimulus in the remaining 80% of trials. A total of 600 stimuli were presented. The task lasted for 20 minutes approximately. No breaks were allowed.

### EEG recording and analysis

EEG data were recorded at 1000 Hz using a 30-channel actiCHamp System (Brain Products GmbH, Munich, Germany) with active electrodes positioned according to the 10-20 EEG International System and referenced to the Cz electrode. The cap was adapted to individual head size, and each electrode was filled with Signa Electro-Gel (Parker Laboratories, Fairfield, NJ) to optimize signal transduction. Participants were instructed to avoid body movements as much as possible, and to keep their gaze on the center of the screen during the task. Electrode impedances were kept below 10 kΩ. EEG preprocessing was conducted using custom Matlab scripts and the EEGLAB [27] and Fieldtrip [28] Matlab toolboxes. EEG data were resampled at 500 Hz, bandpass filtered offline from 1 and 40 Hz to remove signal drifts and line noise, and re-referenced to a common average reference. Horizontal electrooculograms (EOG) were recorded by bipolar external electrodes for the offline detection of ocular artifacts. Independent component analysis was used to detect and remove EEG components reflecting eye blinks [29].

*Spectral power analysis*. Electrodes presenting abnormal power spectrum were identified via visual inspection and replaced by spherical interpolation. Processed EEG data from each experimental period (Warm-up, Exercise, Cool Down) were subsequently segmented to 1-s epochs. The spectral decomposition of each epoch was computed using Fast Fourier Transformation (FFT) applying a symmetric Hamming window and the obtained power values were averaged across experimental periods.

*ERSP analysis*. Task-evoked spectral EEG activity was assessed by computing ERSP in epochs extending from –500 ms to 500 ms time-locked to stimulus onset for frequencies between 4 and 40 Hz. Spectral decomposition was performed using sinusoidal wavelets with 3 cycles at the lowest frequency and increasing by a factor of 0.8 with increasing frequency. Power values were normalized with respect to a –50 ms to 0 ms pre-stimulus baseline and transformed into the decibel scale.

### Statistical analysis

Spectral power main effects of Session (light intensity, Moderate-to-high intensity) were separately tested for significance at each period (Warm-up, Exercise, Cool Down). In the absence of strong a priori hypotheses, we used a stepwise, cluster-based, non-parametric permutation test [30] (Fieldtrip toolbox) without prior assumptions on any frequency range or area of interest. The algorithm performed a t-test for dependent samples on all individual electrodes x frequencies pairs and clustered samples with positive and negative t-values that exceeded a threshold (p < 0.05, two-tailed) based on spatial and spectral adjacency. These comparisons were performed for each frequency bin of 1Hz and for each electrode. Cluster-level statistics were then calculated by taking the sum of the t-values within each cluster. The trials from the two datasets (light intensity, Moderate-to-high intensity) were randomly shuffled and the maximum cluster-level statistic for these new shuffled datasets was calculated. The above procedure was repeated 5000 times to estimate the distribution of maximal cluster-level statistics obtained by chance. The proportion of random partitions that resulted in a larger test statistic than the original one determined the two-tailed Monte-Carlo p-value.

In addition, ERSP main effects of Condition (light intensity, Moderate-to-high intensity) for each stimulus (target 1, target 2 and standard) were also analysed by applying the cluster-based permutation test. In order to reduce the possibility that the type II error rate was inflated by multiple comparisons correction, we grouped data into four frequency bands: Theta (4–8 Hz), Alpha (8–14 Hz), lower Beta (14–20 Hz) and upper Beta 1 (20-40 Hz). Note that the time window of interest in target trials was restricted to the first 300 ms after target onset in order to avoid an overlap with behavioural responses based on average reaction time (RT). The time window of interest for standard trials was fixed to the first 500 ms after the stimulus onset.

Behavioural data from both experiments were analysed using a within-participants factor of Session (light intensity, Moderate-to-high intensity) for RT and accuracy (ACC) as dependent variables. All analyses were completed using statistical non-parametric permutation tests with a Monte Carlo approach [31][32].

## Results

### Spectral power analysis

The analysis of tonic spectral power showed a significant main effect of Session for the exercise period (all *ps* < .02). Two positive clusters (frequency-localization) were found and were statistically significant: one global cluster (25 electrodes) in low frequencies (1-5 Hz), cluster *p* < .001, and one parieto-occipital cluster (11 electrodes) in fast frequencies (8-40 Hz), cluster *p* = .017. The analysis revealed an overall increase in the power of frequencies during the moderate-to-high intensity exercise period in comparison to light intensity (see Fig. 1). There were no statistically significant between-session differences in warm-up and cool down periods (all cluster *ps* ≥ .1).

**Figure 1.**
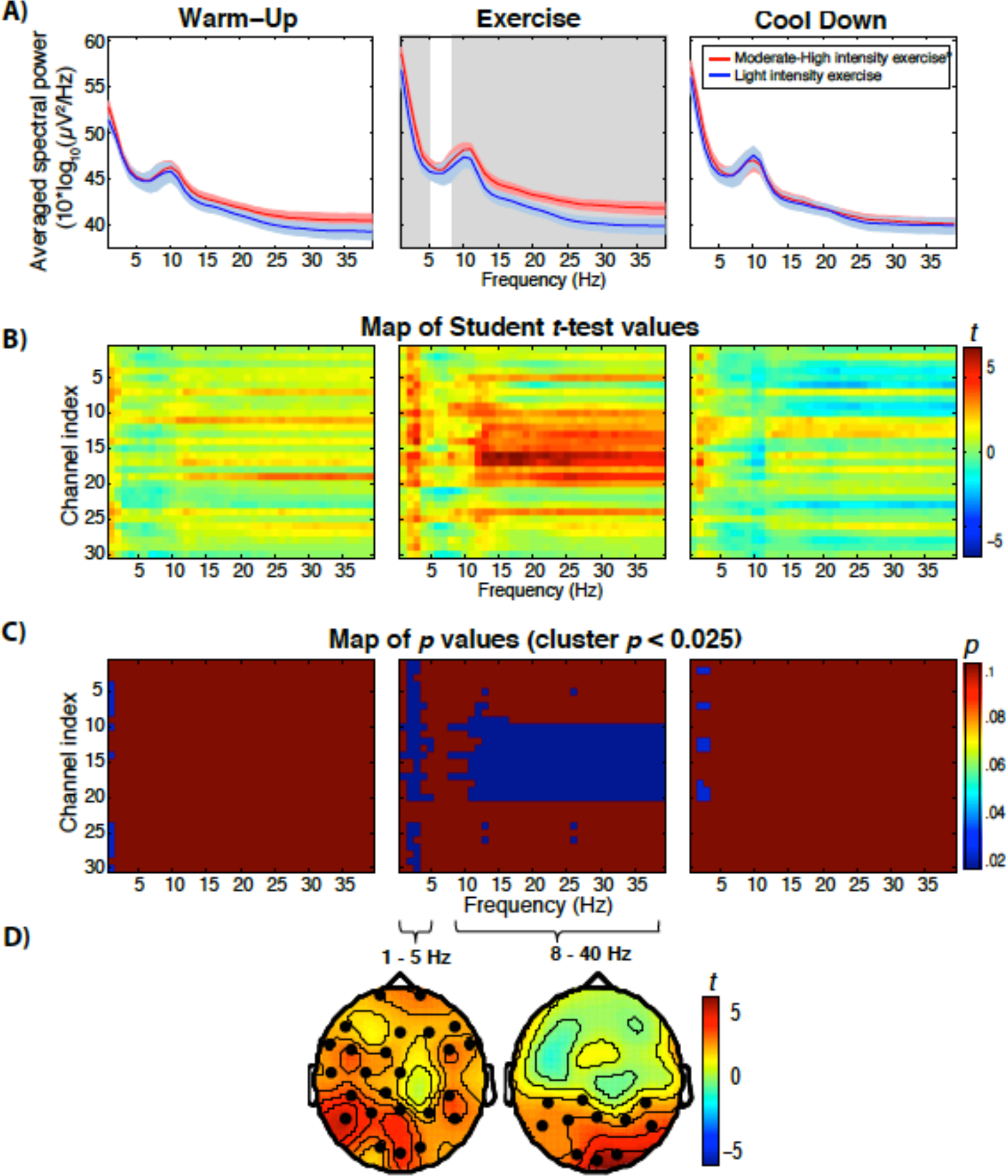
Modulation of brain power spectrum as a function of exercise intensity. (A) Differences in the averaged EEG power spectrum across subjects between moderate-to-high intensity (red lines) and light intensity (blue lines) exercise at the three experimental periods (warm-up, exercise and cool down). Red and blue shaded areas represent 95% confidence intervals. Statistically significant differences are marked by grey area (B) Parametric paired t-test maps comparing the relative power across frequency bands (x-axes) and channels (y-axes) during moderate-to-high intensity and light intensity exercise at warm-up, exercise and cool down (blue: decreases; red: increases). (C) Each image illustrates the statistical significance (*p* values) of the *t*-maps depicting only the significant clusters with p < 0.025. (D) Topographies depict t-test distribution in all electrodes, showing the spatial characteristics of the increase in power of low frequencies across the whole surface localization during moderate-to-high exercise, and the increase in high frequencies in parieto-occipital areas during moderate-to-high exercise. No significant between-intensity differences were found at warm-up and cool down.

### ERSP analysis

Fig. 2 shows the time-locked oscillatory activity of the oddball task during both exercise periods. The analysis of the ERSP revealed a significant main effect of session for standard trials in alpha and lower beta bands (all *ps* < .015). The alpha band analysis revealed only an occipital positive cluster (4 electrodes) between 150-500 ms after the onset of the standard, *p* = .014, with higher alpha spectral power during moderate-to-high intensity exercise compared to light intensity (see Fig 3a). The lower Beta band analysis showed only a parieto-occipital significant positive cluster (8 electrodes) between 300-500 ms after the onset of the standard, *p* = .009. Lower Beta frequency band exhibited a higher spectral power in the moderate-to-high intensity session than in the light intensity session (see Fig 3b).

**Figure 2.**
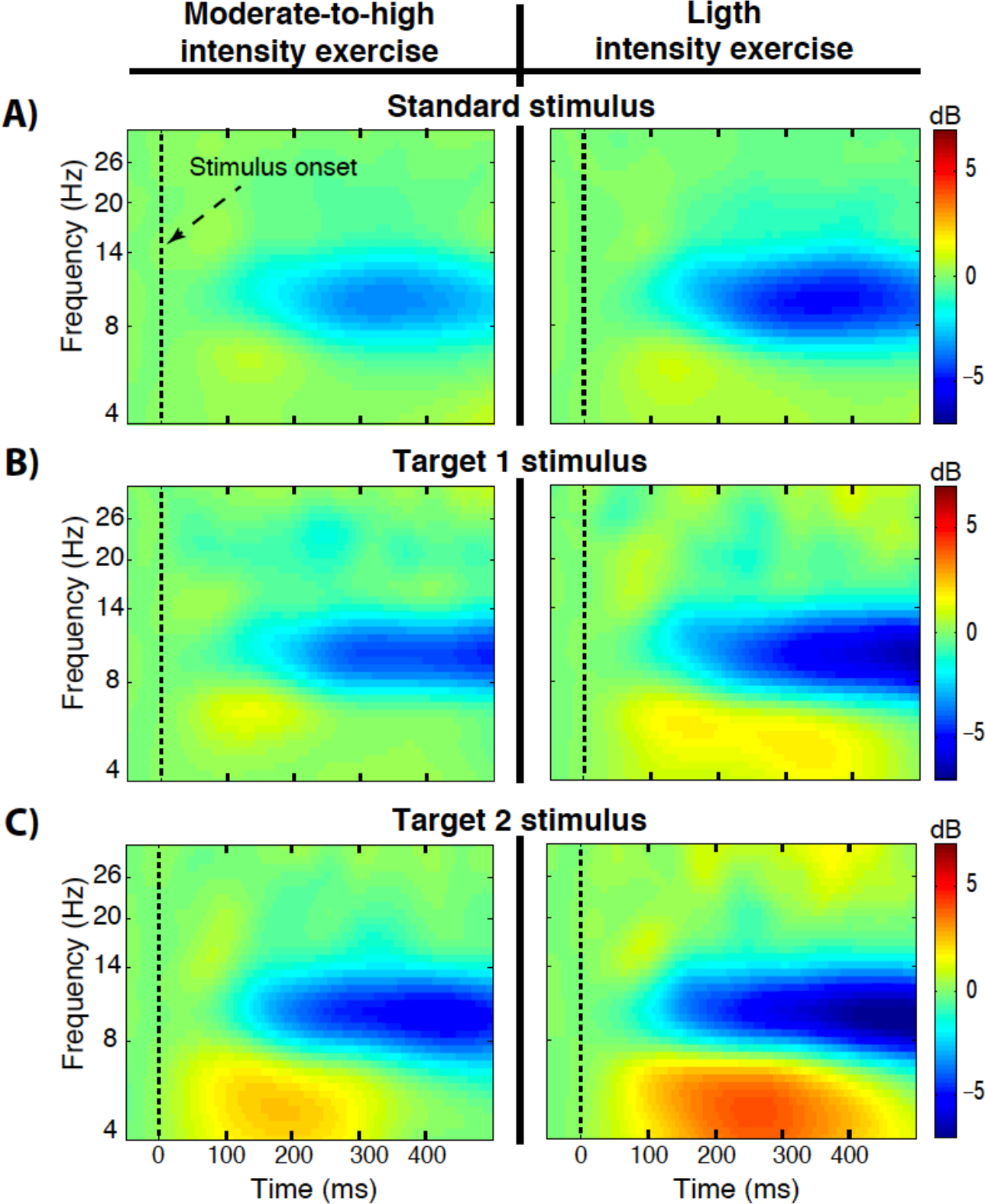
Event-related spectral perturbation of oddball task during exercise. Time-locked spectral power averaged over the occipital channels (Oz, O1 and O2) during moderate-to-high intensity (left column) and light intensity (right column) exercise for all stimuli (standard, target 1 and target 2). Each panel illustrates time-frequency power across time (x-axes) and frequency (y-axes) during moderate-intensity and light-intensity exercise (blue: decreases; red: increases).

The target 2 trials analysis revealed a significant main effect of Session in the theta band (*p* < .01). Two negative clusters (time-localization) were found. Only the largest global negative cluster (7 electrodes) between 190-250 ms after the onset of the target was significant, cluster *p* = .004. Target 2 trials (red squares) during moderate-to-high intensity exercise session evoked lesser theta power than during light intensity exercise condition (see Fig 3c). The analysis of the other frequency bands for target 1, target 2 and standard trials did not yield significant effects (all *ps* ≥. 05).

**Figure 3.**
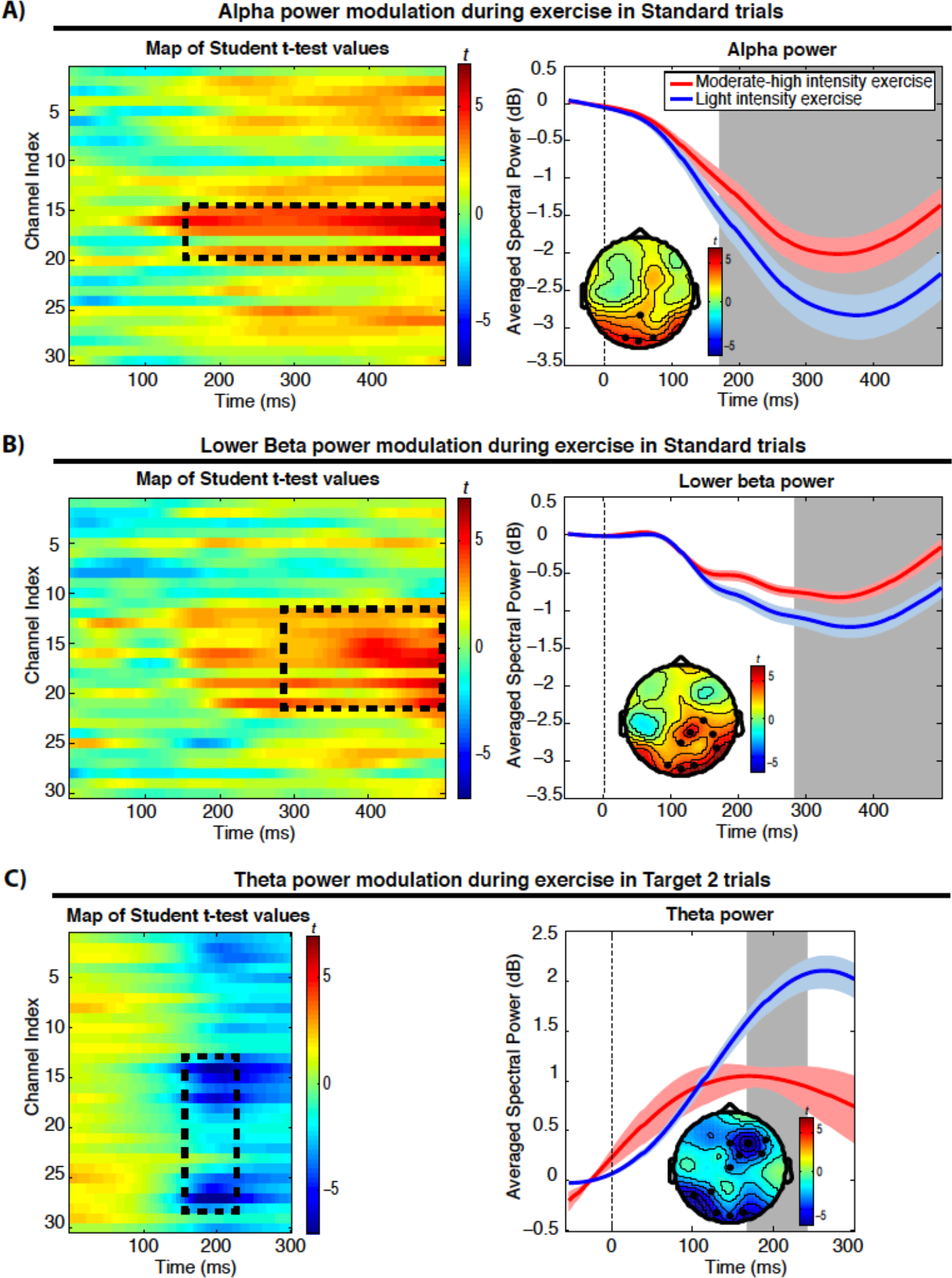
Event-related spectral perturbation significant main effects of session. (A) Alpha frequency band [8-14 Hz] parametric paired t-test maps comparing the averaged spectral power across subjects over time (x-axes) and frequency (y-axes) during moderate-to-high intensity and light intensity exercise in standard trials. The enclosed areas denote significant clusters of channels and time with p < 0. 025. Right panel shows alpha power across time at the occipital-parietal cluster (4 electrodes) in standard trials. The shaded area represents the latency range where significant differences between exercise sessions were found. Red and blue shaded areas represent 95% confidence intervals. The topography depicts t-test distribution across surface localization, showing the spatial characteristics of the higher power suppression of alpha in parieto-occipital electrodes during moderate-to-high exercise. (B) Parametric paired t-test maps comparison between moderate-to-high exercise and light exercise in lower beta frequency band [14-20 Hz] to standard trials. Right panel shows lower beta power across time at the parieto-occipital cluster (8 electrodes) in standard trials. (C) Theta frequency band [4-8 Hz] parametric paired t-test maps comparing moderate-to-high intensity and light intensity exercise in target 2 trials (red squares). Right panel shows theta power across time at the globally-localized cluster (7 electrodes) in target 2 trials. Note that the time window of interest in target trials was restricted to the first 300 ms in order to avoid neural activity overlapping with behavioural responses. The analysis of the other frequency bands for standard, target 1, and target 2 stimuli did not yield significant effects.

### Behavioural Performance

Table 1 provides mean and 95% confidence intervals of behavioural measures as a function of session. Analysis of response RTs and ACC did not reveal statistically significant differences between sessions, (all *ps* > .05).

**Table 1.**
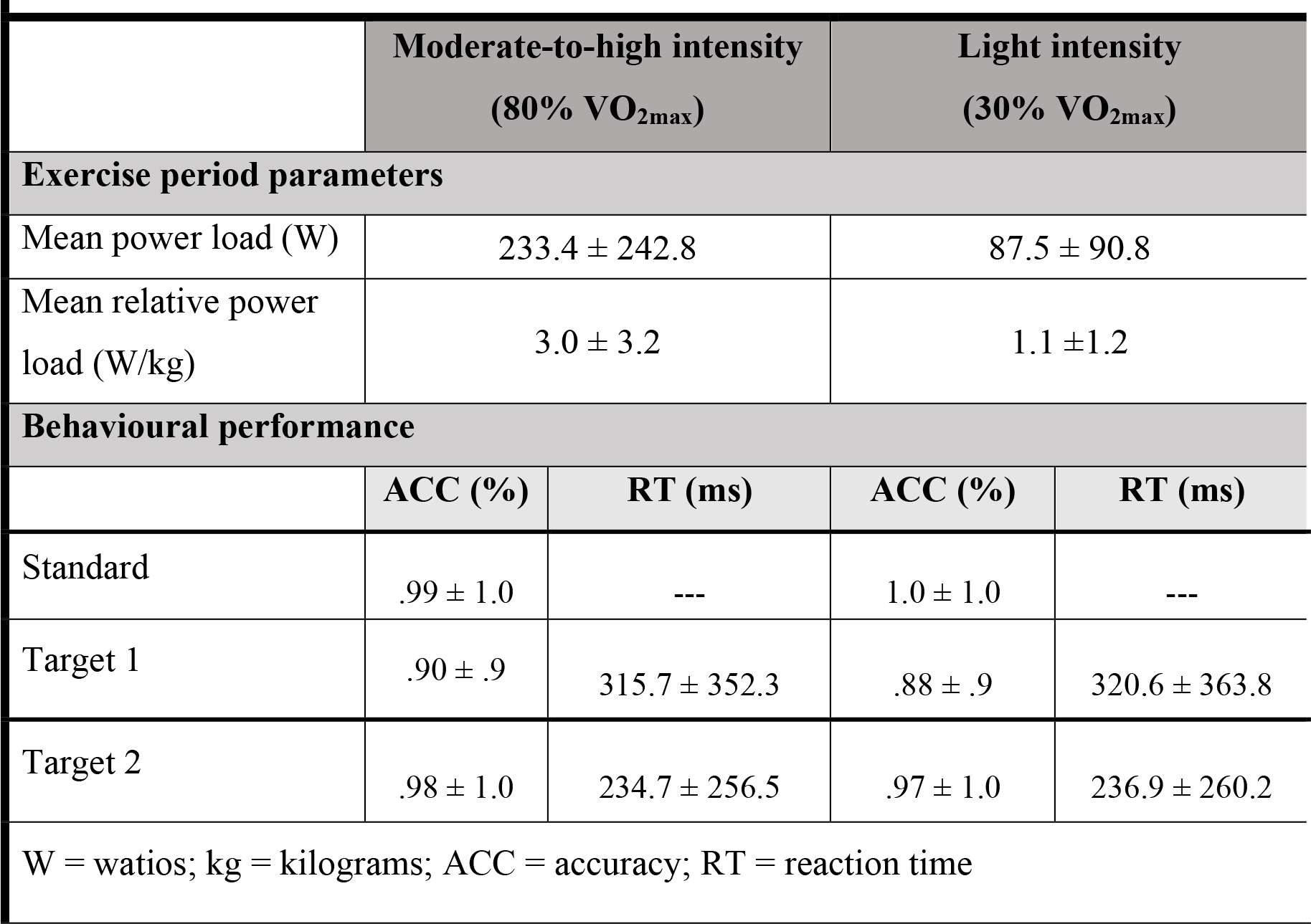
Mean and 95% confidence intervals of descriptive exercise-intensity parameters and behavioural performance for the moderate-to-high intensity and low intensity sessions.

## Discussion

In the present study, we investigated the oscillatory brain activity of young adults, during a single bout of aerobic exercise (cycling) at 80% of maximum aerobic capacity compared to a light intensity exercise (control) session while performing a visual oddball task. We found that acute exercise at moderate-to-high intensity induced changes in oscillatory brain activity at the tonic and transient (event-related) level with respect to light intensity.

The overall power increase across the frequency spectrum during moderate-to-high intensity exercise with respect to light intensity is in line with the results of the only meta-analysis to date that has addressed this issue [12]. Indeed, Crabbe and Dishman [12] found no evidence of the selective effect of exercise on alpha frequency band at frontal localizations suggested by previous empirical research [9][10][11][33]. Interestingly, the between-intensity differences were unspecific of surface localization in slow frequencies, while in faster frequencies the differences arose from parieto-occipital sites. These results partially contradict previous studies that have shown changes in oscillatory brain activity during exercise localized in anterior sites [9][34].

Several studies have suggested that the theta band is selectively enhanced by the presentation of novel stimuli, linking it to the orienting responses associated with novelty processing [35][36]. Moreover, alpha activity suppression has been associated with cognitive engagement to the task [37][38]during oddball paradigms similar to the one uses in the present study [39][40]. Thus, the present theta and alpha results might represent an attentional resources regulation during stimulus processing in order to maintain optimal task performance. In line with this interpretation, and given that task behavioural performance was not significantly different between sessions, one could argue that the reduced theta power to target 2 trials (the more salient stimuli) and the lower suppression of alpha and lower beta bands during the moderate-to-high intensity exercise session with respect to the light intensity session, was a sign of brain efficiency. Results from previous ERP studies during a single bout of exercise have also pointed to processing efficiency during moderate-to-high intensity exercise [17][41] [42]. At this point, it is important to note that both exercise sessions were matched in terms of dual-task demands. Participants were instructed to maintain the pedaling cadence constant between 60 and 90 rpm in both sessions, therefore any variation in brain functioning was due to the physiological changes induced by the particular exercise intensity.

Exercising elicits a wide set of physiological changes such as increases in core temperature, cortical blood flow, heart rate, and catecholamine concentration [43] which have generally been recognized as a potential mechanism underlying the acute exercise effect on brain function. Interestingly, we found a higher global increase of oscillatory brain activity during the moderate-to-high intensity session than the light intensity session. This latter result is consistent with recent accounts that have linked acute exercise to enhanced activation/arousal (that relates to the overall activation/excitability of cortical neurons [44][45]). However, the ERSP results in our study suggest that the effect of acute exercise cannot be explained as a mere overall increase of oscillatory brain activity but to a specific way of brain functioning during exercise (at moderate-to-high intensity).

To conclude, the present study contributes to further understanding of how the brain works during acute exercise and the underlying neural mechanisms of the positive relationship between physical exercise, brain function and cognition in young adults. Importantly, a profound knowledge of the factors that support this beneficial relationship is especially critical for public health, as a sedentary lifestyle has been linked to a series of chronic diseases and to a deterioration of cognitive function and performance in many daily life activities.

## Acknowledgments

This work was supported by the “Ministerío de Economia y Competitividad” (grant numbers PSI2013-46385-P and PSI2016-75956-P) and the “Junta de Andalucía” (grant number SEJ-6414). P.Ch.I. acknowledges support from W. M. Keck Foundation, and the Office of Naval Research (ONR Grant No. 000141010078). The funders had no role in study design, data collection and analysis, decision to publish, or preparation of the manuscript. We thank to all the participants who took part in the experiment.

